# Wnt/Fgf crosstalk is required for the specification of tracheal basal progenitor cells

**DOI:** 10.1101/405159

**Authors:** Zhili Hou, Qi Wu, Xin Sun, Huaiyong Chen, Yu Li, Yongchun Zhang, Munemasa Mori, Ying Yang, Ming Jiang, Jianwen Que

## Abstract

Basal progenitor cells are critical for the establishment and maintenance of the tracheal epithelium. However, it remains unclear how these progenitor cells are specified during foregut development. Here, we found that ablation of the Wnt chaperon protein Gpr177 (also known as Wntless) in the epithelium causes significant reduction in the numbers of basal progenitor cells accompanied by cartilage loss in *Shh-Cre;Gpr177^loxp/loxp^* mutants. Consistent with the association between cartilage and basal cell development, Nkx2.1^+^p63^+^ basal cells are co-present with cartilage nodules in *Shh-Cre;Ctnnb1^DM/loxp^* mutants which keep partial cell-cell adhesion but not the transcription regulation function of ß-catenin. More importantly, deletion of *Ctnnb1* in the mesenchyme leads to the loss of basal cells and cartilage concomitant with the reduced transcript levels of Fgf10 in *Dermo1-Cre;Ctnnb1^loxp/loxp^* mutants. Furthermore, deletion of *Fgf receptor 2* (*Fgfr2*) in the epithelium also leads to significantly reduced numbers of basal cells, supporting the importance of the Wnt/Fgf crosstalk in early tracheal development.

## Introduction

Basal cells are multipotent progenitor cells responsible for the generation of the airway epithelium during development and injury-repair (Hong et al., 2004; Que et al., 2009; Rock et al., 2009; Yang et al., 2018). Previous studies indicate that the epithelial-mesenchymal interactions are critical for basal cell development although the underlying molecular mechanism remains largely unknown (Hines et al., 2013; Volckaert et al., 2013). It has been shown that the numbers of basal cells are positively correlated with the differentiation of mesenchymal cells into cartilage at the early stage of tracheal development (Hines et al., 2013). Prior to the separation of the trachea from the anterior foregut, the growth factor Fgf10 is enriched in the ventral mesenchyme where cartilage progenitor cells arise (Que et al., 2007). Interestingly, ubiquitous Fgf10 overexpression promotes basal cell lineage commitment while suppressing ciliated cell differentiation (Volckaert et al., 2013). In addition, ubiquitous overexpression of the Wnt inhibitor Dkk1 at E10.5 but not E12.5 also leads to the increased numbers of basal cells (Volckaert et al., 2013). Wnt signaling is critical for initial specification of respiratory cells from the early foregut. Loss of Wnt2/2b which are enriched in the ventral foregut mesenchyme results in failed specification of respiratory progenitor cells (Nkx2.1^+^) (Goss et al., 2009). Consistently, deletion of the canonical Wnt signaling mediator ß-catenin also leads to lung and tracheal agenesis, and the anterior foregut becomes an esophageal-like tube lined with stratified squamous epithelium underlined by extensive basal progenitor cells (Goss et al., 2009; Harris-Johnson et al., 2009).

We and others previously showed that the respiratory cell fate is specified properly despite severe vasculature abnormalities following deletion of the Wnt chaperon protein Gpr177 (also known as Wntless) in *Shh-Cre;Gpr177^loxp/loxp^* mutants. Interestingly, in this study we found a significant loss of basal progenitor and cartilage cells in these mutants. Deletion of *Ctnnb1* encoding ß-catenin in the mesenchyme also leads to the loss of basal progenitor cells and cartilage concomitant with the reduced levels of Fgf10 in the trachea of *Dermo1-Cre;Ctnnb1^loxp/loxp^* mutants. Moreover, the numbers of basal progenitor cells are also significantly reduced when the Fgf10 receptor *Fgfr2* is deleted in the epithelium. Together these findings support that in the developing trachea epithelial Wnts activate ß-catenin in the mesenchyme to modulate Fgf10 levels which in turn regulate basal cell specification through epithelial Fgfr2.

## Results and Discussion

### Blocking Wnt secretion from the epithelium leads to a reduced number of basal progenitor cells and cartilage defects in the trachea of *Shh-Cre;Gpr177^loxp/loxp^* mutants

We previously showed that deletion of *Gpr177* in the epithelium results in the abnormal differentiation and proliferation of vascular smooth muscle cells in the developing lung, and the mutants succumb at birth due to severe pulmonary hemorrhage (Jiang et al., 2013). A recent study demonstrated that deletion of *Gpr177* also leads to tracheal cartilage defects in these mutants (Snowball et al., 2015). We asked whether basal cell specification is affected upon *Gpr177* deletion given the correlation of cartilage and basal cell numbers (Hines et al., 2013). Consistent with previous findings (Snowball et al., 2015), *Gpr177* deletion results in the loss of cartilage progenitor cells (Sox9^+^) while smooth muscle cells (SMA^+^) are expanded in the trachea of *Shh-Cre;Gpr177^loxp/loxp^* mutants (Fig. 1A-B). Intriguingly, basal cells (p63^+^) are rarely detected in the trachea of *Shh-Cre;Gpr177^loxp/loxp^* mutants at different developing stages examined (Fig. 1A-B, n=3 for each stage). These results suggest that Wnts from the epithelium act in both autocrine and paracrine manners to regulate tracheal development. Notably, *Gpr177* deletion did not to affect the specification of respiratory cells from the early foregut and all the epithelial cells express Nkx2.1 (Fig. 1A-B). Increased differentiation of ciliated (α-acetylated tubulin^+^) but not club cells (Cc10^+^) was also observed in the tracheal epithelium at late stage (Fig. 1B). In addition, although loss of *Gpr177* leads to the reduced proliferation of both epithelial and mesenchymal cells, the difference between mutants and wildtype controls is not significant (Fig. 1C, p>0.05, n=3 for each).

**Fig. 1.**
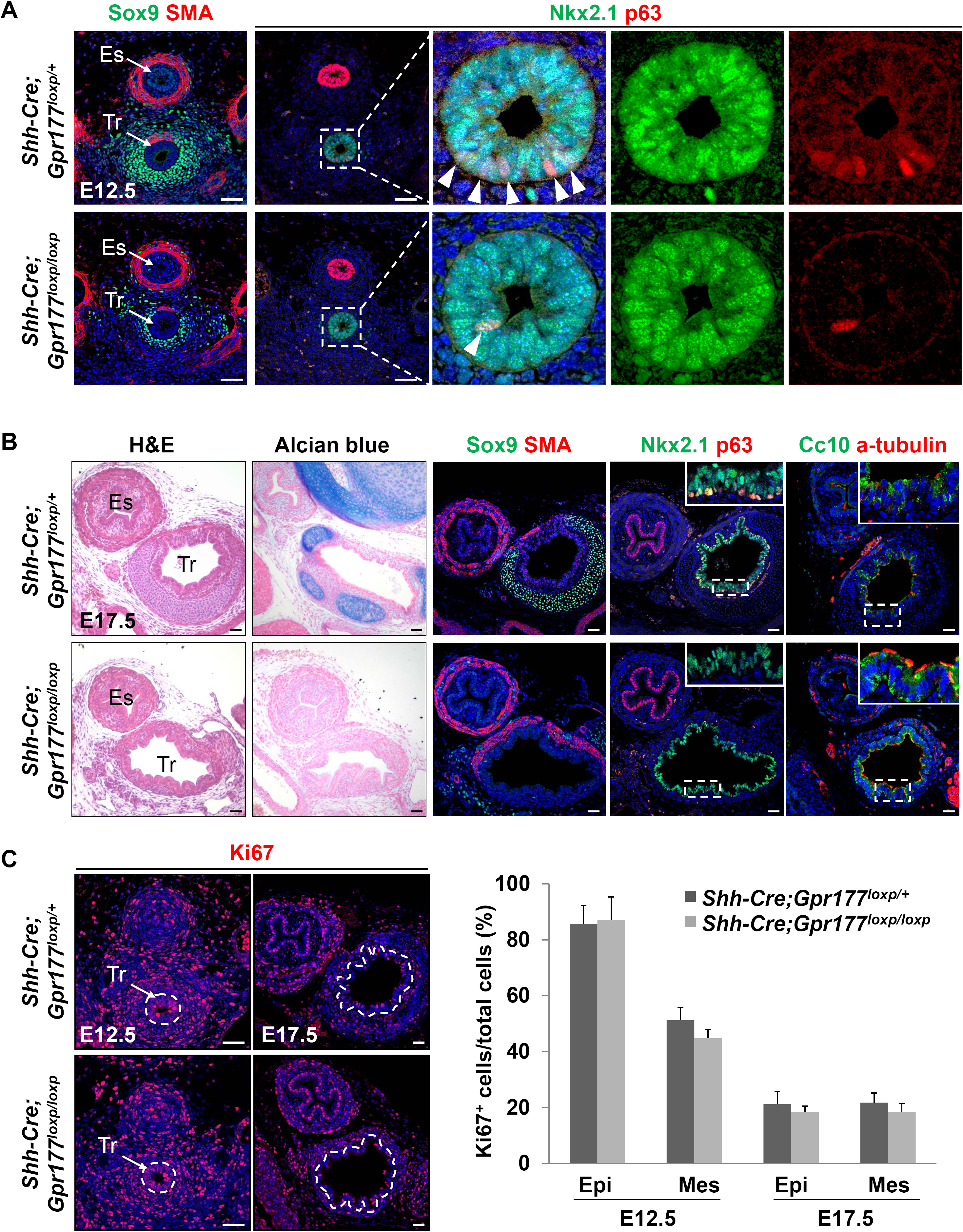
Loss of epithelial Wnt secretion leads to abnormal development of tracheal cartilage and basal cell. **(A)** Loss of epithelial *Gpr177* causes dramatic reduction in the numbers of cartilage progenitor cells (Sox9^+^) and basal cells (p63^+^) in the trachea of *Shh-Cre; Gpr177^loxp/loxp^* mutant at E12.5. **(B)** Loss of epithelial *Gpr177* leads to the loss of cartilage (Alcian Blue^+^ Sox9^+^) and reduction in the number of basal cells at E17.5. Note the increased number of ciliated cells in the mutant trachea. **(C)** The proliferation of both epithelial and mesenchymal cells is slightly but not significantly decreased in the mutant trachea (p>0.05 n=3 for each). Data are represented as mean ± SEM. Abbreviation: Es: esophagus; Tr: trachea. Scale Bar: 50 μm.

### Presence of tracheal basal cells (Nkx2.1^+^p63^+^) and cartilage nodules in the unseparated foregut of *Shh-Cre; Ctnnb1^DM/loxp^* mutants

β-catenin has two major roles, mediating Wnt-activated transcription regulation and cell-cell adhesion functions (Heuberger and Birchmeier, 2010). Thus far genetic studies assessing the role of ß-catenin in the developing lung have relied on the *Ctnnb1^loxp^* allele which ablates both transcription regulation and cell-cell adhesion functions upon Cre-mediated recombination (Brault et al., 2001; De Langhe et al., 2008; Goss et al., 2009; Stenman et al., 2008). Although many of the phenotypic changes seem to recapitulate what have been observed in mutants lacking Wnt ligands (Goss et al., 2009; Stenman et al., 2008), it is unclear whether the cell-cell adhesive function of β-catenin contributes to lung and tracheal development.

Another mouse line containing a mutated *Ctnnb1* allele (*Ctnnb1* double mutant; hereafter referred as *Ctnnb1^DM^*) was recently established (Valenta et al., 2011). This mutant form of β- catenin includes a single amino acid change (aspartic acid mutated to Alanine, D164A) in the first Armadillo repeat of β-catenin, which prevents TCF-dependent transcription regulation while maintaining the ability to mediate cellular adhesion (Valenta et al., 2011). Notably, *Ctnnb1^DM/DM^* mutants die at E10.5 (Gay et al., 2015). We combined this allele with the *Ctnnb1^loxp^* allele to address whether ß-catenin acts in the epithelium to regulate basal cell development. As expected, β-catenin protein is retained in the epithelial junction of *Shh-Cre;Ctnnb1^DM/loxp^* but not *Shh-Cre;Ctnnb1^loxp/loxp^* mutants (Fig. S1). Similar to *Shh-Cre;Ctnnb1^loxp/loxp^* mutants, the anterior foregut remains a single-lumen tube in *Shh-Cre;Ctnnb1^DM/loxp^* embryos (Fig. 2A). Consistent with previous findings (Goss et al., 2009; Harris-Johnson et al., 2009), the unseparated foregut is specified as an esophageal-like muscular tube lined by squamous basal cells (Nkx2.1^-^p63^+^) in *Shh-Cre;Ctnnb1^loxp/loxp^* mutants (Fig. 2B). By contrast, the ventral side of the unseparated foregut in *Shh-Cre;Ctnnb1^DM/loxp^* mutants demonstrates respiratory cell differentiation (Nkx2.1^+^) underlined by cartilage nodules (Sox9^+^) (Fig. 2C). These ventral epithelial cells express the columnar cell marker Krt8, but not the squamous cell marker Krt13 (Fig. S2B). More importantly, tracheal basal cells (Nkx2.1^+^p63^+^) are present in the proximity of the cartilage nodules, confirming the association of cartilage and basal cell development (Hines et al., 2013). Of note is that some residual ciliated and club cells are also present in the ventral foregut of *Shh-Cre;Ctnnb1^DM/loxp^* mutants (Fig. S2C). Taken together these results suggest that the cellular adhesion function of ß-catenin plays roles in tracheal development. That being said, we can’t rule out the possibility that some residual ß-catenin mediated transcription regulation activities remain present in *Shh-Cre;Ctnnb1^DM/loxp^* mutants, even though TCF-transactivation dependent Wnt signaling is ablated in various studies using the *Ctnnb1^DM^* mouse line (Azim et al., 2014; Gay et al., 2015; Valenta et al., 2016; Valenta et al., 2011).

**Fig. 2.**
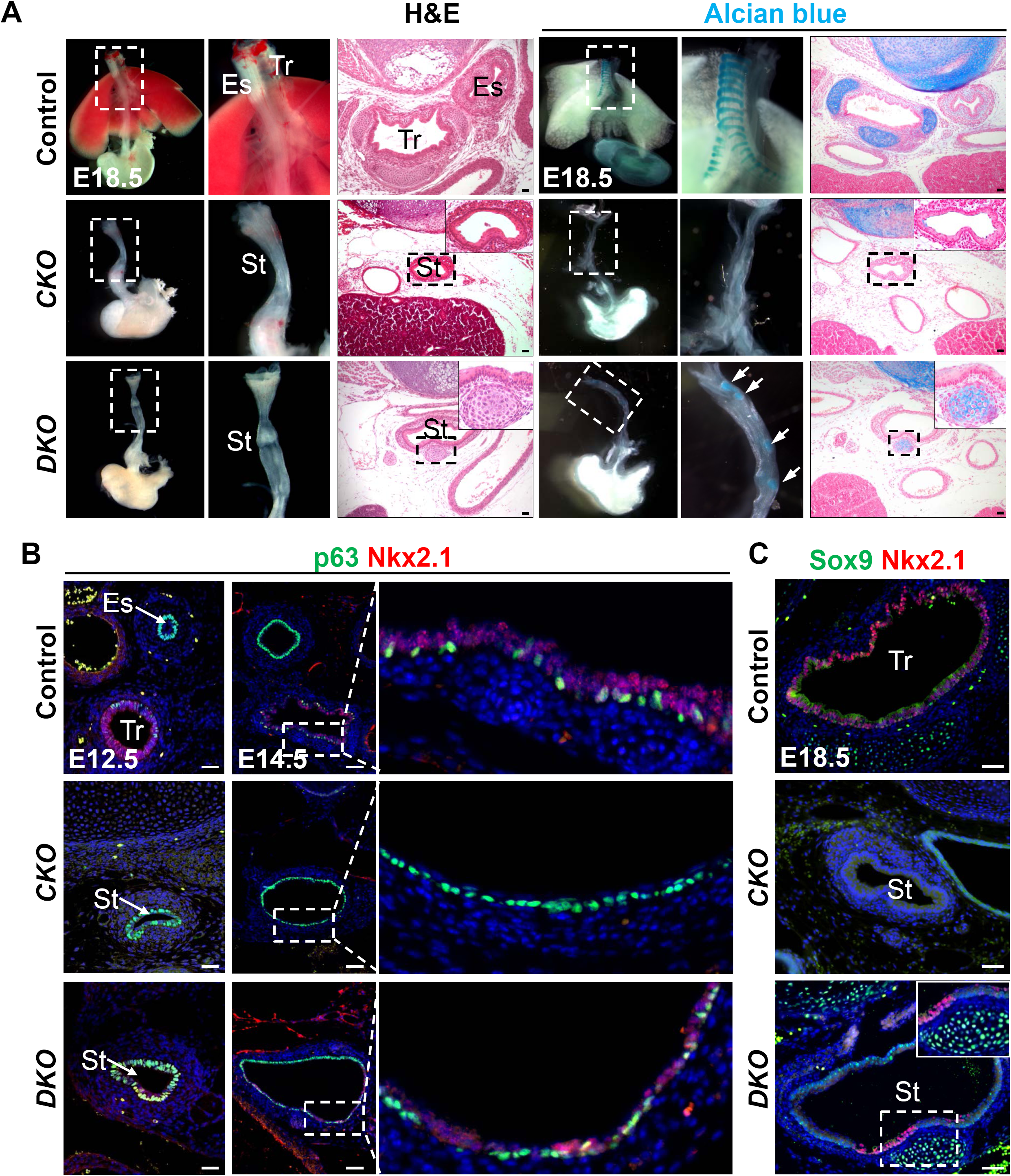
Residual tracheal basal cells and cartilage nodules in *Shh-Cre; Gpr177^DM/loxp^* mutants. **(A)** Failed separation of the trachea and esophagus in *Shh-Cre; Gpr177^loxp/loxp^* (*Ctnnb1^CKO^*) and *Shh-Cre; Gpr177^DM/loxp^* (*Ctnnb1^DKO^*) mutants. Note the stratified squamous epithelium lining the muscular tube in *Ctnnb1^CKO^* mutants and residual cartilage nodules (Alcian blue^+^) in *Ctnnb1^DKO^* mutants. **(B)** Tracheal basal cells (Nkx2.1^+^ p63+) are present in the ventral side of the unseparated foregut in *Ctnnb1^DKO^* but not *Ctnnb1^CKO^* mutants. **(C)** Residual respiratory epithelium (Nkx2.1^+^) is underlined by cartilage nodules (Sox9^+^) in *Ctnnb1^DKO^* mutants. Abbreviation: Es, esophagus; Tr, Trachea; St, single lumen tube. Scale bar: 50 μm.

### Epithelial Wnts regulate basal cell and cartilage development through mesenchymal ß- catenin

Loss of epithelial Wnts inhibits the development of tracheal cartilage and basal cells in *Shh-Cre; Gpr177^loxp/loxp^* mutants. We asked whether epithelial Wnts directly regulate basal cell and cartilage development through β-catenin in the mesenchyme. We deleted *β-catenin* with *Dermo1-Cre* which is active in the mesenchymal progenitor cells as early as E10.5 (Hines et al., 2013; Sala et al., 2011). As previously described, the trachea is separated from the early foregut but is shortened, accompanied by simplified lung branching morphogenesis in *Dermo1-Cre; Ctnnb1^loxp/loxp^* mutants (De Langhe et al., 2008). Interestingly, cartilage progenitor cells (Sox9^+^) are absent in the trachea of mutants at E11.5 (Fig. 3A). Sox9 immunostaining further confirmed the loss of cartilage at E14.5 (Fig. 3A). By contrast, smooth muscle cells (SMA^+^) extend to the ventral side of the trachea (Fig. 3A). Loss of cartilage is accompanied by significant reduction in the number of basal progenitor cells which were barely detected in the trachea at E11.5 (n=3) and E14.5 (n=5) (Fig. 3B). Notably, p63^+^ cells are also barely detected in the ventral foregut epithelium prior to the separation of the trachea from the foregut (Fig. 3B). Together these data support that β-catenin in the mesenchyme is a critical mediator for epithelial Wnts to regulate cartilage and basal cell specification in the developing trachea.

**Fig. 3.**
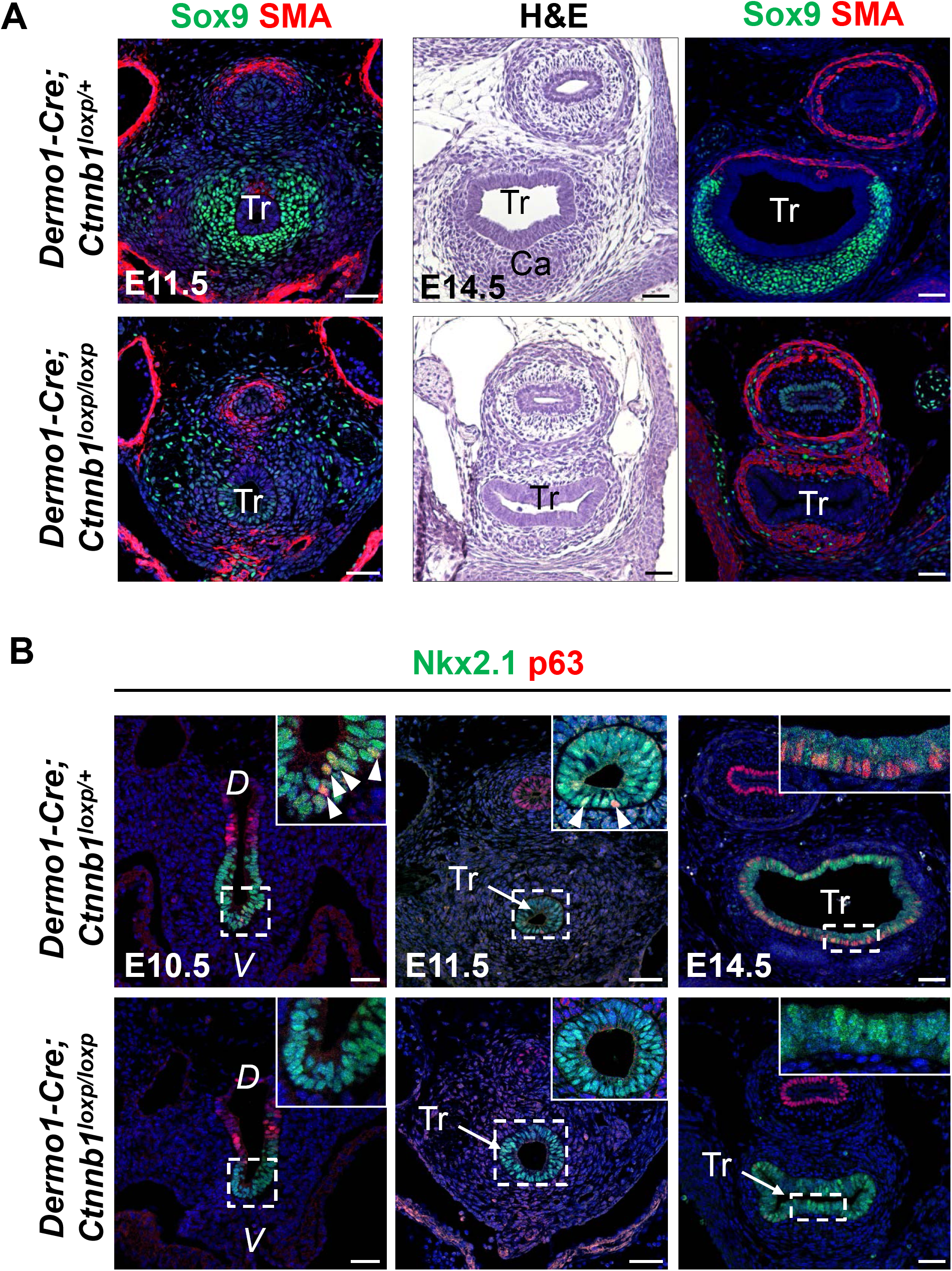
Deletion of *Ctnnb1* in the mesenchyme causes the loss of tracheal cartilage and basal cells in *Dermo1-Cre; Ctnnb1^loxp/loxp^* mutants. **(A)** Tracheal cartilage is absent in *Dermo1-Cre; Ctnnb1^loxp/loxp^* mutants. Note some neuronal-like cells (Sox9^+^) in the mutant trachea. **(B)** Basal cells are barely detected in both ventral and dorsal sides of the mutant trachea at different developmental stages. Abbreviation: Es, esophagus; Tr, trachea; Ca, cartilage; *D*, dorsal; *V*, ventral. Scale bar: 50 μm.

### Mesenchymal ß-catenin regulates basal cell specification through crosstalk with Fgf10/Fgfr2 signaling

Previous studies have shown that Fgf10 overexpression leads to an increased number of basal cells in the airways (Volckaert et al., 2013). We therefore asked whether the transcript levels of Fgf10 are downregulated in *Dermo1-Cre;Ctnnb1^loxp/loxp^* mutants at E11.5. Consistent with mitigated Wnt activities, the transcript levels of the Wnt/β-catenin downstream targets Axin2 and Lef1 are significantly decreased (Fig. 4A). Interestingly, we also observed dramatical reduction in the transcript levels of Fgf10 upon *Ctnnb1* deletion in the mesenchyme (Fig. 4A). Previously we and others have shown that Fgf10 is expressed in the ventral mesenchyme of the foregut prior to tracheal-esophageal separation and then restricted to the inter-cartilage compartment after tracheal cartilage condensation occurs (Que et al., 2007; Sala et al., 2011). By contrast, the Fgf10 receptor Fgfr2 is uniformly expressed in the epithelium (Sala et al., 2011). We hypothesized that decreased Fgf signaling contributes to the loss of basal cells in *Dermo1-Cre;Ctnnb1^loxp/loxp^* mutants. We therefore deleted *Fgfr2* in the early foregut epithelium using *Shh-Cre*. Consistent with previous finding, loss of *Fgfr2* leads to lung agenesis and truncated trachea (Sala et al., 2011). Conditional loss of *Fgfr2* also leads to less condensed cartilage although the alternative pattern of smooth muscle and cartilage seems not perturbed (Fig. 4B). Importantly, similar to what has been observed in *Dermo1-Cre;Ctnnb1^loxp/loxp^* mutants, the numbers of basal cells are significantly reduced in the trachea of *Shh-Cre;Fgfr2^loxp/loxp^* mutants (Fig. 4C). These findings support a model whereby Fgf10 from the mesenchyme under the control of ß-catenin is critical for the specification of basal cells in the developing trachea. A crosstalk between Hippo signaling and Fgf10 has been shown to regulate basal cell-fueled epithelial regeneration in the adult trachea (Volckaert et al., 2017). Here, our findings support that the ß-catenin/Fgf10/Fgfr2 axis plays an important role in basal cell specification during early tracheal development.

**Fig. 4.**
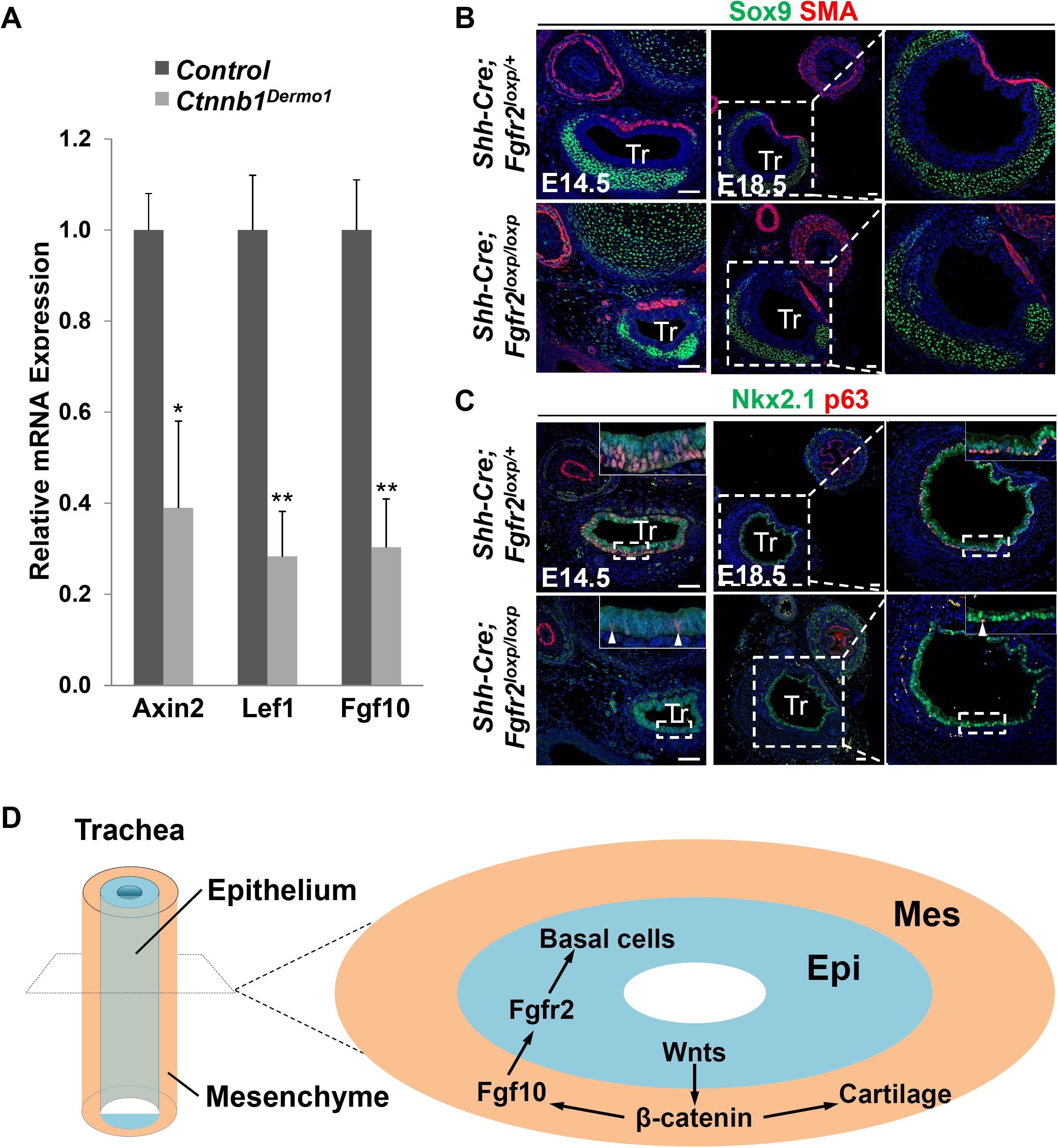
Mesenchymal β-catenin regulates the specification of basal cell through Fgf10/Fgfr2 signaling. **(A)** The reduced transcript levels of the Wnt/β-catenin downstream targets Axin2 and Lef1 and Fgf10 in the trachea of *Dermo1-Cre; Ctnnb1^loxp/loxp^* mutants (\**p*<0.05; \*\**p*<0.01; *n*=3 independent experiments). **(B)** Deletion of *Fgfr2* leads to less-condensed cartilage in the trachea of *Shh-Cre;Fgfr2^loxp/loxp^* mutants. **(C)** Loss of *Fgfr2* causes significant reduction in the number of basal cells (arrowheads). **(D)** Diagram depicts that the epithelial-mesenchymal interaction mediated by Wnt/β-catenin signaling regulates tracheal basal cell and cartilage development. Epithelial Wnt proteins directly activate mesenchymal β-catenin to promote the expression of Fgf10 which in turn activates epithelial Fgfr2 to modulate basal cell specification. Abbreviation: Tr: trachea; Epi: epithelium; Mes: mesenchyme. Scale bar: 50 μm.

In summary, our study revealed that Wnt proteins secreted from the respiratory epithelium is critical for both tracheal cartilage and basal cell development. We found residual ß- catenin, presumably mediating cell-cell adhesion function is critical for the specification of tracheal epithelium and cartilage development in *Shh-Cre;Ctnnb1^DM/loxp^* mutants. On the other hand, ß-catenin in the mesenchyme plays significant roles in the specification of basal progenitor cells and cartilage. Our further genetic studies suggest that mesenchymal ß-catenin regulates Fgf10 which relays to its receptor Fgfr2 in the epithelium to regulate basal cell specification (Fig. 4D).

## Acknowledgement

We are grateful that Dr. Konrad Basler (University of Zurich) shared with us the *Ctnnb1^DM^* mouse line. This work is partly supported by R01HL132996 and the Price Family Foundation.

## Author contributions

M.J. and J.Q. designed experiments, analyzed data, and wrote the manuscript. Z.H. and M.J. performed the experiments. M.M. provided *Shh-Cre;Fgfr2^loxp/loxp^* embryos. Q.W., X.S., H.C., Y.Z. Y.L., and Y.Y. assisted with mouse genetics.

## Figure Legends

**Fig. S1.**
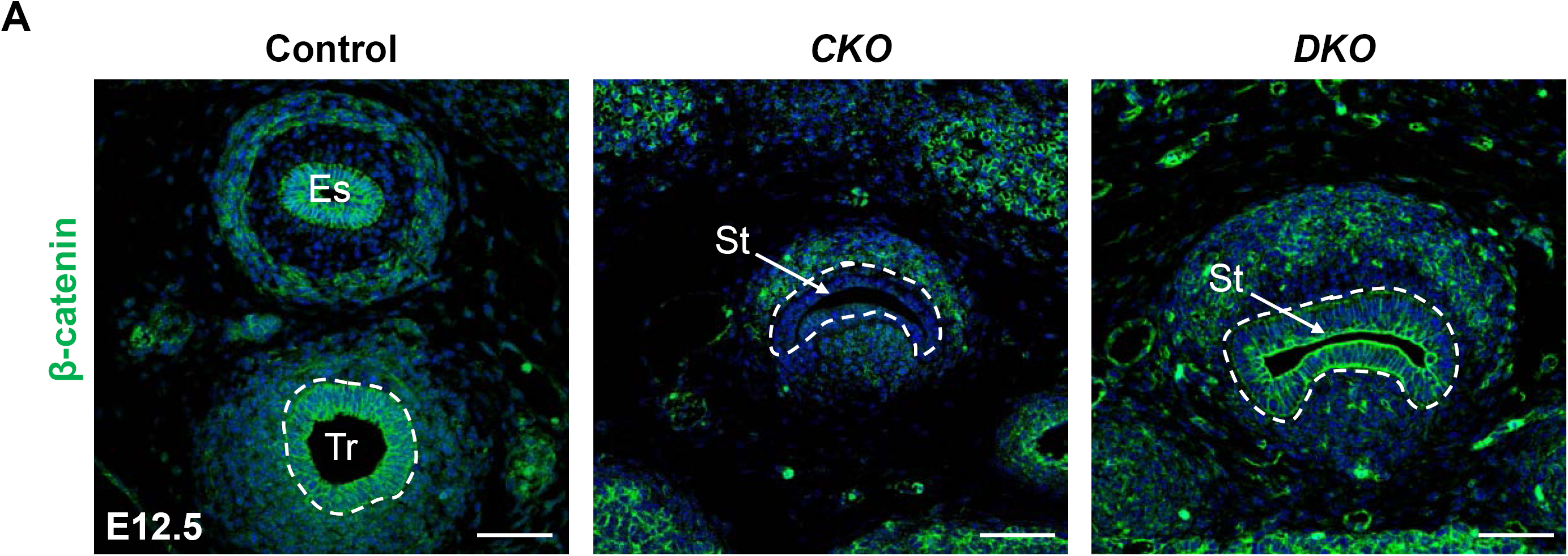
β-catenin protein is retained in the epithelial junction of *Ctnnb1^DKO^* but not *Ctnnb1^CKO^* mutants. Abbreviation: Es: esophagus; Tr: trachea; St: single lumen tube; *CKO: Shh-Cre;Ctnnb1^loxp/loxp^; DKO: Shh-Cre;Ctnnb1^DM/loxp^*. Scale bar: 50 μm.

**Fig. S2.**
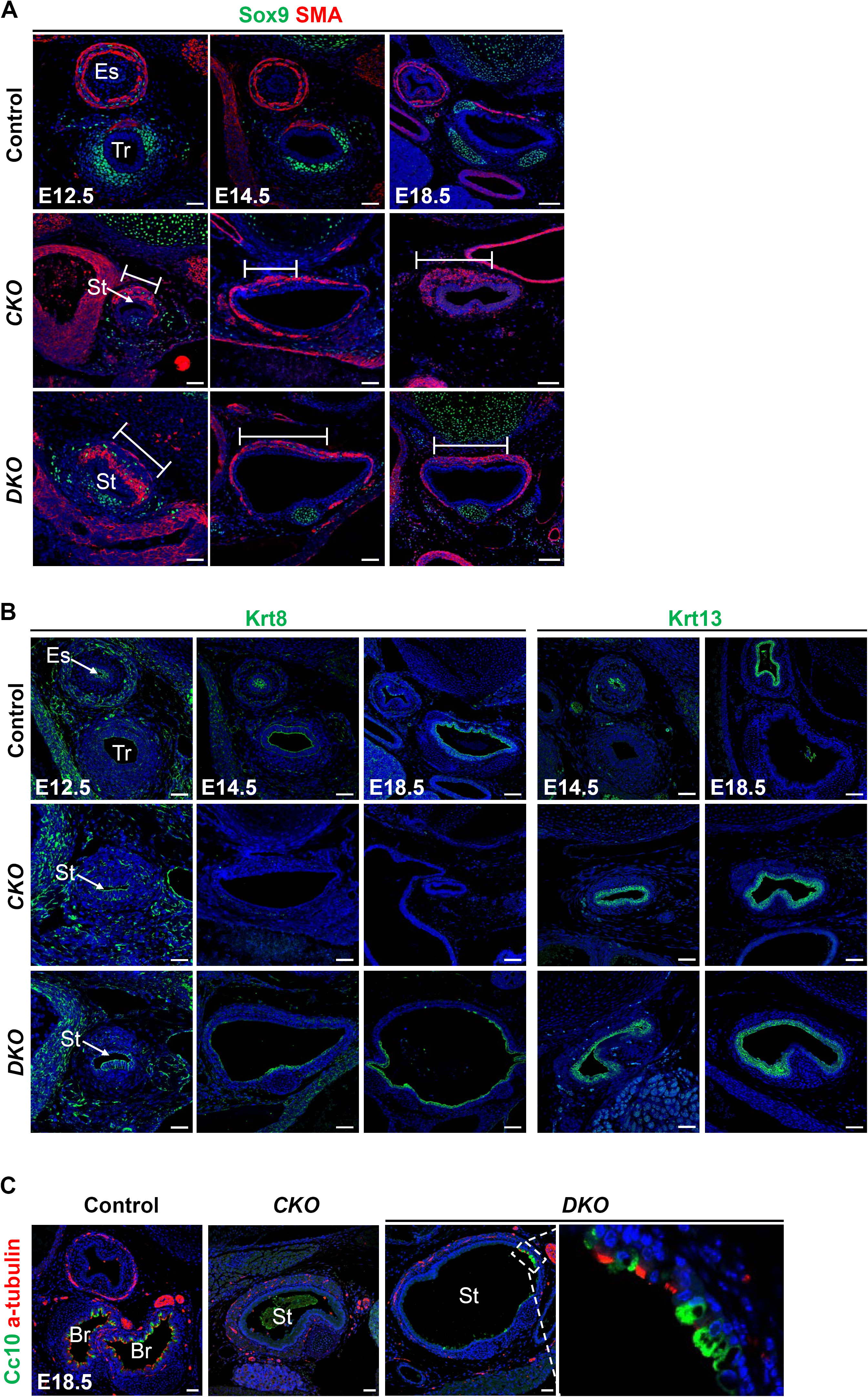
Presence of residual respiratory epithelium in the unseparated foregut tube in *Ctnnb1^DKO^* but not *Ctnnb1^CKO^* mutants. **(A)** Presence of cartilage nodules in *Ctnnb1^DKO^* but not *Ctnnb1^CKO^* mutants. Note expanded smooth muscle cells in the foregut of both *Ctnnb1^CKO^* and *Ctnnb1^DKO^* mutants. **(B)** The residual respiratory epithelial cells in *Ctnnb1^DKO^* mutants express the columnar cell marker Krt8, but not the squamous cell protein Krt13 which is enriched throughout the esophagus-like tube in *Ctnnb1^CKO^* mutants. **(C)** Presence of club cells (Cc10^+^) and ciliated cells (a-tubulin^+^) in the ventral side of the unseparated foregut of *Ctnnb1^DKO^* mutants at E18.5. Abbreviation: Es, esophagus; Tr, trachea; Br, bronchus; St, single lumen tube; *CKO: Shh-Cre;Ctnnb1^loxp/loxp^; DKO: Shh-Cre;Ctnnb1^DM/loxp^*. Scale bar: 50 μm.

## Materials and Methods

### Mice

The *Shh-Cre*, *Dermo1-Cre*, *Gpr177^loxp/loxp^*, *Ctnnb1^loxp/loxp^* and *Fgfr2^loxp/loxp^* mouse strains, and genotyping methods have been reported previously (Brault et al., 2001; Fu et al., 2009; Harfe et al., 2004; Yu et al., 2003). *Ctnnb1^DM/+^* mice were kindly provided by Dr. Konrad Basler of University of Zurich (Valenta et al., 2011). All mice were maintained in the University’s animal facility according to institutional guidelines. All mouse experiments were conducted in accordance with procedures approved by the Institutional Animal Care and Use Committee.

### Tissue processing, histology and immunostaining

For paraffin sections, tissues were fixed in 4% paraformaldehyde overnight and processed as previously described (Jiang et al., 2017). For cryo-sections, tissues were fixed in 4% paraformaldehyde in PBS at 4°C overnight, placed in 30% sucrose in PBS, and embedded in OCT. The primary antibodies used for immunostaining analysis include: rabbit anti-Nkx2.1 (1:500, ab76013, Abcam); mouse anti-p63 (1:500, CM163, Biocare); rabbit anti-Sox9 (1:1000, AB5535, Millipore); mouse anti-smooth muscle actin (SMA) (1:2000, A2547, Sigma); chicken anti–KRT8 (1:1000, ab107115, Abcam); rabbit anti-Krt13 (1:1000, ab92551, Abcam); rabbit anti-β-catenin (1:200, 8480S, Cell Signaling Technology); rabbit anti-Cc10 (1:500, 06-263, Millipore); mouse anti-α-acetylated tubulin (1:5000, T7451, Sigma); mouse anti-Ki67 (1:500, 550609, BD Biosciences). Fluorescent secondary antibodies were used for detection and visualization. Images were obtained using Nikon SMZ1500 Inverted microscope (Nikon). Confocal images were obtained with a Zeiss LSM T-PMT confocal laser-scanning microscope (Carl Zeiss).

### Alcian blue staining

Whole lungs were dissected in PBS solution and fixed in 95% EtOH. Alcian blue staining was performed as previously described (Jiang et al., 2017). Briefly, whole lungs and sections were treated with 3% acetic acid solution for 3 minutes, then stained in Alcian blue (A3157, Sigma) for 5 minutes and counterstained with Nuclear Fast Red (N8002, Sigma).

### Reverse transcription and real-time PCR

RNA extraction and reverse transcription was performed using the Super-Script III First-Strand SuperMix (Invitrogen) according to the manufacturer’s instructions. cDNA was subjected to quantitative real-time PCR using the StepOnePlus Real-Time PCR Detection System (Applied Biosystems) and iTaq Universal SYBR Green Supermix (Bio-Rad). All real-time quantitative PCR experiments were performed in triplicate. The prime sequences were as follows: β-actin forward 5’-CGGCCAGGTCATCACTATTGGCAAC-3’ and reverse 5’- GCCACAGGATTCCATACCCAA-3’; Axin2 forward 5’-CAGCCCTTGTGGTTCAAGCT and reverse 5’-GGTAGATTCCTGATGGCCGTAGT-3’; Lef1 forward 5’- GCAGCTATCAACCAGATCC-3’ and 5’-GATGTAGGCAGCTGTCATTC-3’; Fgf10 forward 5’-CGGGACCAAGAATGAAGACT-3’ and reverse 5’-AGTTGCTGTTGATGGCTTTG-3’.

### Statistical analysis

Statistical analysis was done by Student’s t-test. Data are presented as mean ± s.e.m.; p values < 0.05 were considered statistically significant.

